# Transcranial Direct Current Stimulation enhances long-term retention after 5 days of lower-limb motor skill learning

**DOI:** 10.64898/2026.02.11.705269

**Authors:** August Lomholt Kvistad, Lasse Jespersen, Jonas Rud Bjørndal, Lasse Christiansen, Anke Ninija Karabanov, Jesper Lundbye-Jensen

**Affiliations:** Movement & Neuroscience, Department of Nutrition, Exercise and Sports, University of Copenhagen, Copenhagen, Denmark; Danish Research Centre for Magnetic Resonance, Department of Radiology and Nuclear Medicine, Copenhagen University Hospital - Amager and Hvidovre, Copenhagen, Denmark, Kettegård Allé 30, 2650 Hvidovre, Denmark; Department of Neuroscience, University of Copenhagen, Blegdamsvej 3B, 2200 Copenhagen N, Denmark

**Author notes:** Correspondence: August Lomholt Kvistad, Movement and Neuroscience, Department of Nutrition, Exercise and Sports, University of Copenhagen, Denmark.

**Keywords:** Transcranial Electric Stimulation, tDCS, Motor skill learning, Bayesian analysis

## Abstract

Transcranial direct current stimulation (tDCS) holds the potential to affect behavior by modulating ongoing neural activity, and tDCS paired with hand motor practice can enhance motor learning. While augmenting the behavioral benefits of motor practice is relevant for neurorehabilitation following central and peripheral lesions to the motor system, as well as in sports, the short- and long-term effects of tDCS targeting the mesial motor cortex (M1-Leg) during lower-limb motor skill practice remain unexplored. We tested whether five days of anodal tDCS over M1-Leg during training of a sequential visuomotor tracking task improves within- and between-session learning and one-week retention. Participants were randomized to skill practice with active tDCS, skill practice with sham stimulation, or volume-matched non-skilled ankle movements with sham stimulation. Changes in corticospinal excitability accompanying skill and nonskill motor practice with real and sham tDCS, were assessed as motor evoked potential amplitudes recorded from the tibialis anterior muscle at rest. Compared to non-skill practice, motor skill practice yielded robust sequence-specific performance gains, which were transferred to the untrained leg and persisted for at least one week after practice ended. Concurrent tDCS did not increase learning within or between training sessions, but it did lead to improved one-week retention compared to sham stimulation. Corticospinal excitability did not increase after practice and was unaffected by tDCS. These findings suggest that combining lower-limb motor skill practice with tDCS over M1-Leg can strengthen retention of skill learning without measurable changes in resting corticospinal excitability. This is relevant for motor practice scheduling in neurorehabilitation.

**Key points:**

- Transcranial direct current stimulation (tDCS) is a weak electrical brain stimulation that may boost learning when paired with motor practice; however, its effects during lower-limb skill training are not well known.
- We tested whether stimulation over the leg area of the motor cortex during 5 days of ankle skill training improves learning, consolidation, and delayed retention.
- Ankle skill training produced clear sequence-specific improvements that transferred to the untrained leg and were still present 1 week later.
- Stimulation did not increase short-term learning, but it did improve 1-week retention compared to sham stimulation.
- Corticospinal excitability assessed based on motor evoked potentials elicited by transcranial magnetic stimulation did not change with motor training or stimulation, suggesting that the observed positive effect of tDCS on delayed retention may arise from other brain network processes relevant to long-term motor learning and memory.

## Introduction

Human motor skill learning encompasses our ability to acquire, refine, and retain new motor skills, underpinning everything from daily activities to high-level athletic performance and rehabilitation after injury. The process of skill learning depends on repeated practice of movements and movement sequences with high spatiotemporal acuity. Plastic changes in the central nervous system are necessary for skill learning, and repeated, goal-directed practice is essential for inducing the synaptic and network-level plasticity that underlies durable skill gains (Peters *et al*., 2017; Roth & Ding, 2024). Transcranial direct current stimulation (tDCS) offers a scalable adjunct to motor practice; it is portable, low-cost, and suitable for at-home use. Recent evidence has shown that pairing motor practice with concurrent tDCS improves motor skill learning (Nielsen et al., 2025). However, most evidence comes from single-session practice of upper-limb movements, rendering both the longevity of tDCS effects as well as the potential for boosting effects of lower extremity motor practice unexplored. Given the prominent role of lower extremity rehabilitation in both progressive neurological disorders and following lesions to the motor system, it is pertinent to clarify whether the application of tDCS during motor practice carries long-lasting benefits for lower extremity motor learning.

Direct current stimulation modulates neuronal activity by inducing weak electric fields (E-fields) that shift firing probabilities, as demonstrated both in vitro and in vivo (Bindman *et al*., 1962, 1964; Reato *et al*., 2010; Rahman *et al*., 2017; Lafon *et al*., 2017). Although tDCS yields E-fields in the targeted brain tissue that are substantially weaker than those needed to drive firing in vitro robustly (Rahman *et al*., 2013; Vöröslakos *et al*., 2018; Liu *et al*., 2018), these intensities are sufficient to bias ongoing neural activity rather than activate quiescent circuits (Bikson & Rahman, 2013). Consequently, tDCS can be considered a non-invasive and safe (Antal *et al*., 2017) neuromodulatory tool that, when paired with task-specific practice, can increase network engagement and amplify plasticity in already active neurons, while leaving unrelated pathways comparatively unaffected (Fritsch *et al*., 2010; Reis & Fritsch, 2011; Madhavan & Shah, 2012; Kronberg *et al*., 2017, 2020; Gellner *et al*., 2020; Wang *et al*., 2022). Therefore, functionally targeted tDCS applied concurrently with motor practice may offer a mechanistic route to behavioral or therapeutic gains by selectively reinforcing networks recruited during practice.

In support of this, findings from predominantly upper limb motor practice lend credence to the notion that anodal tDCS applied to the primary motor cortex (M1) reinforces practice-induced plasticity, thereby enhancing motor skill learning (Buch et al., 2017; Nielsen, et al., 2025; Reis et al., 2009). Evidence in favor of lower-limb applications is contrastingly sparse. A few single-session studies suggest that anodal tDCS concurrent with practice can enhance within-session learning and 24-hour retention (Shah *et al*., 2013; Sriraman *et al*., 2014; Foerster *et al*., 2018); however, whether repeated pairing across days accumulates benefits for lower-limb long-term skill learning remains unknown.

Here, we address this gap by applying anodal M1-tDCS during five days of lower-limb motor skill sequence training and assessing within- and between-session learning (online/offline learning) and one-week retention of skill learning. Specifically, we employ a parallel three-arm design, including a group engaged in motor practice with concurrent active tDCS (SKILL-STIM), one group undergoing motor practice with sham (SKILL-SHAM), and a control group that practices without a fixed sequence and without demands for precision but with an equivalent number and amplitude of movements combined with sham tDCS (CON). To separate sequence-specific gains from general improvements in motor acuity, we included a near-transfer condition using a randomized target sequence. To test practice-dependent corticospinal modulation over time, we additionally investigate corticospinal excitability (CSE) via motor-evoked potentials (MEPs) before and after practice on Days 1 and 5 (Christiansen *et al*., 2018). We used Bayesian predictive modelling to test a priori graded hypotheses for both immediate learning and one-week retention (SKILL-SHAM > CON; SKILL-STIM > SKILL-SHAM). In parallel, we evaluated whether skilled training modulates corticospinal excitability (CSE), operationalized as the pre-to-post change in resting MEP amplitudes from the tibialis anterior muscle. Specifically, we tested whether skill practice yields larger CSE increases than non-skill control (SKILL-SHAM > CON) and whether concurrent anodal tDCS further potentiates this change (SKILL-STIM > SKILL-SHAM), consistent with state-dependent facilitation of plasticity mechanisms during task engagement.

## Methods

### Ethical Approval

The study was reviewed and approved by the local ethics committee (protocol H-17019671), and all experimental procedures were conducted in accordance with the *Declaration of Helsinki*. The study was not preregistered in a database before recruitment. All participants received written and oral information about the study and provided written and verbal informed consent prior to enrolment.

### Participants

Sixty healthy and able-bodied participants (28 females, aged 20-30 years) were enrolled in this study. The inclusion criteria were verified using a standardized health screening questionnaire confirming no history of neurological, psychiatric, or musculoskeletal disorders and no current medication use. One participant withdrew during the first experimental session. The participants were randomized into three groups: (1) motor skill practice and concurrent tDCS (SKILL-STIM), (2) motor skill practice and sham tDCS (SKILL-SHAM), and (3) non-skill motor practice and sham tDCS (Control; CON). All participants performed motor practice using their non-dominant leg, determined by preferred leg to kick a ball with which aligns with the Waterloo Footedness Questionnaire-Revised (WFQ-R) (Van Melick *et al*., 2017). The participant characteristics are presented in Table 1.

**Table 1:** Participant characteristics. Values are presented as mean ± SD unless otherwise noted; categorical variables are presented as counts. Groups: SKILL-STIM (motor skill practice + real tDCS), SKILL-SHAM (motor skill practice + sham tDCS), CON (non-skill control + sham). Sex (F/M) = female/male counts. Age (years). Height (cm). Weight (kg). MVC dorsiflexion (Nm) = maximal voluntary contraction torque of the ankle dorsiflexors. Stim guess (Real/Sham) = participants’ end-of-study guess of the stimulation condition. Dominant leg (n=R) = number reporting right leg dominance. rMT (% MSO) = resting motor threshold expressed as a percentage of maximum stimulator output. Sleep during the five days (h) = average overnight sleep duration across the five practice days.

| Participant Characteristics | SKILL-STIM | SKILL-SHAM | CON |
| --- | --- | --- | --- |
| Sex (F/M) | 10/10 | 9/11 | 9/11 |
| Age (Years) | 23.9±2.1 | 24.3±3.7 | 24.5±3.3 |
| Height (cm) | 175±9.6 | 178±11.6 | 176.8±7.6 |
| Weight (kg) | 71.9±11.3 | 75,8±13,6 | 70,3±10,3 |
| MVC dorsiflexion (Nm) | 20.8±5.5 | 23.8±6.6 | 21.8±5.2 |
| Stim guess (Real/Sham) | 7/13 | 8/12 | 5/14 |
| Dominant leg n=R | 16 | 20 | 18 |
| rMT, % MSO | 58.5±10.1 | 60±10.8 | 64.7±13.5 |
| Sleep – during the 5 days (hours per night) | 7.7±0.6 | 7.7±0.7 | 7.3±0.9 |

### Experimental design and procedure

A double-blinded randomized between-group design was employed. Both participants and experimenters were blinded to the type of stimulation (real vs. sham tDCS), but not to the type of motor practice (motor skill practice vs. non-skilled movements). Participants were randomly allocated to three groups, stratified by age and sex, prior to the start of the intervention (SKILL-STIM, n = 20; SKILL-SHAM, n = 20; CON, n = 20). The testing and training spanned five consecutive days. SKILL-STIM group received real/active tDCS concurrently with motor skill practice, the SKILL-SHAM group received sham tDCS concurrently with motor skill practice, and the CON group received sham tDCS and performed non-skilled ankle dorsal flexion. The main experiment involved behavioral and electrophysiological testing procedures and practice of the visuomotor task for approximately 20 min/day for 5 consecutive days. All participants returned after seven days for a long-term retention test of skill performance. Before and after the practice sessions on days 1 and 5, corticospinal excitability and maximal muscle compound action potential (M_max_) were measured. Motor practice continued on Days 2–5, with each session comprising six blocks of 12 movement sequences (i.e., trials).

On Day 5, an EEG recording session was inserted prior to the final practice block (no stimulation), which added one extra block (seven blocks in total that day). To maintain the consistency of the behavioral/TMS outcomes across days and to keep the focus of the present report on behavioral and TMS endpoints, EEG data (and analyses tied to the EEG session) are not included here and are presented elsewhere. A randomized block (near transfer task) was added after the final sequence block. The transfer leg performance was assessed on Days 1 and 5. Follow-up testing was conducted on Day 12 to assess skill retention. The experimental design is shown in Fig. 1B, and the procedures for behavioral and electrophysiological testing are described in detail below.

**Figure 1:**
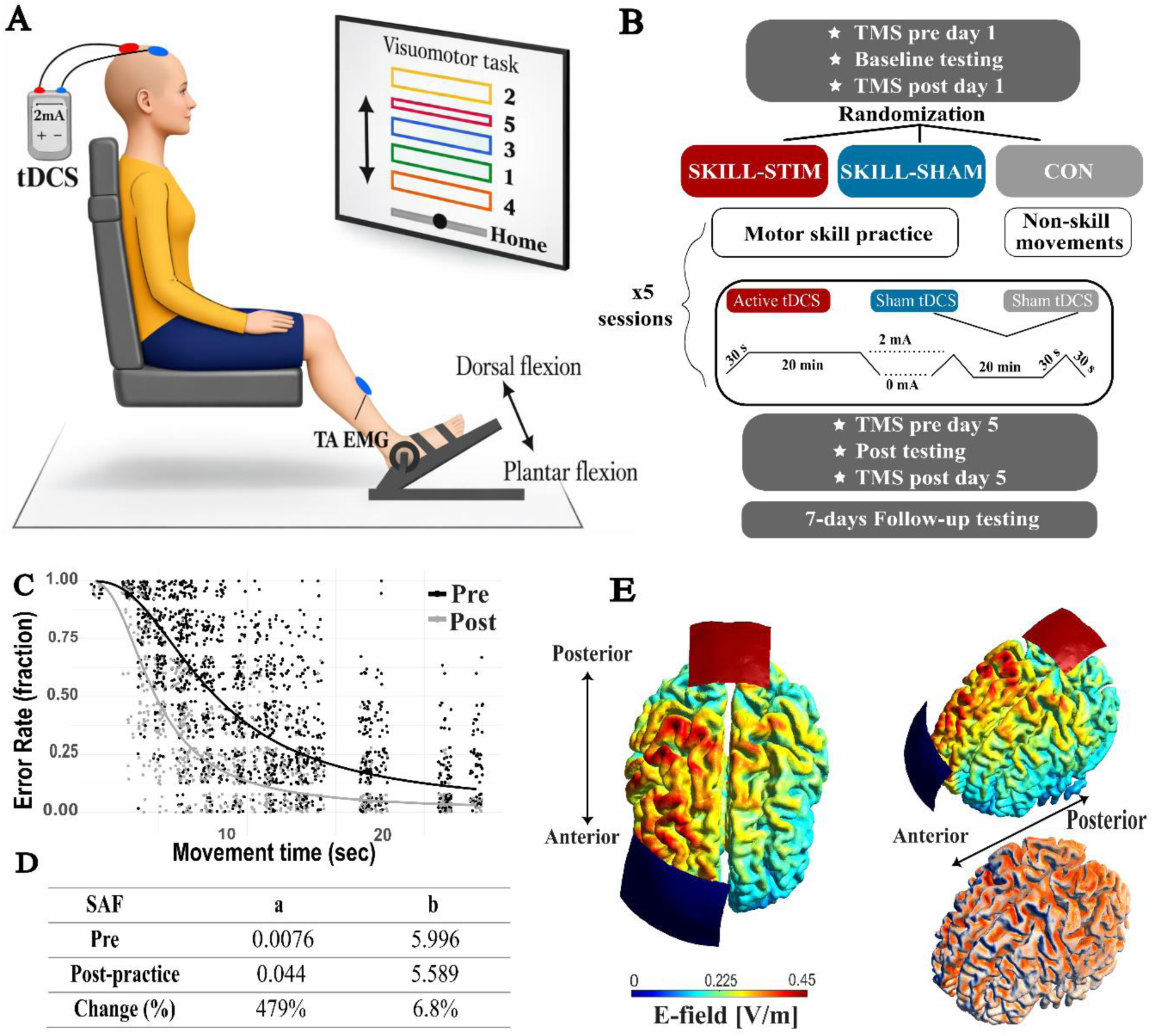
Experimental setup and time course of the experiment. The experimental setup **(A)** and design **(B)** illustrate the visuomotor task and time course of the experiment, respectively. Validation of speed-accuracy function **(C)** in the experimental design involving the lower limb and dynamic dorsal flexion. **D)** Changes in parameters a and b (free parameters) in the speed-accuracy function (SAF). **E)** Computational simulation using SimNIBS4.0 (SimNibs, DRCMR and DTU, Copenhagen and Lyngby, Denmark) of the electric field (EF) induced in the cortex (red corresponds to higher EF (V/m)) and the electrode placement with anode (red) placed close to the vertex and cathode (blue) placed at the supraorbital region contralateral to the performing lower limb. Right lower corner: Inflow of the electric field (orange) and outflow (blue).

### Sequential Visuomotor Dynamic Ankle Flexion Task

The motor learning paradigm was adapted from the well-established Sequential Visual Isometric Pinch Task (SVIPT) (Reis *et al*., 2009) but reconfigured to emphasize dynamic position control of the ankle rather than isometric force control. Therefore, we developed a sequential visuomotor dynamic ankle flexion task. Participants were seated with their non-dominant foot secured in a custom-built dynamic ankle force and position pedal (Fig. 1A), which permitted free dorsiflexion and plantar flexion of the ankle joint. A goniometer tracked the ankle angle, beginning from a standardized resting position of 120°. Each participant’s active range of motion was measured beforehand, and the task movement span was set to 80% of the maximum dorsiflexion excursion. Following presentation of a ‘GO’ cue, participants executed a smooth dorsiflexion through the sequence of five targets (Home–1–Home–2–Home–3–Home–4–Home–5) (Fig. 1A). The ankle angle was mapped in real-time to the vertical cursor position on a computer display. Upon reaching target 5, a 4-s rest period followed. One trial was defined as one complete Home-to-5 sequence; blocks consisted of 12 consecutive trials, followed by a rest interval. The objective was to traverse the entire sequence as rapidly and accurately as possible, and the completion time (CT) was recorded from the movement onset until the cursor reached target 5. Because the visuomotor task is a continuous, multi-segment task, accuracy was quantified as the mean deviation rate per block, that is, the average fraction of gate-crossing errors (either overshoots or undershoots when attempting to hit each target) across all trials, rather than simply the proportion of fully successful trials (Reis et al., 2009). Furthermore, the use of the Tibialis Anterior (TA) muscle enabled the investigation of CSE changes in a single lower limb muscle, which is readily accessible with TMS. The experimental setup is shown in Figure 1A. The visuomotor task source code is available at GitHub (https://github.com/MovementAndNeuroscience/TrackIt-LiteV2).

### Motor practice

A motor practice session consisted of six blocks of 12 trials with a 2-min break between each block. For SKILL-STIM and SKILL-SHAM, the visuomotor task was performed with full visual feedback from the CT and ER in every trial. Intertrial breaks were set to 4 s with a visual countdown. CON group performed non-skill ankle dorsiflexion exercises matched in number to the skilled practice. At trial onset, a baseline target appeared, and its disappearance (screen blackout for 1 s) cued a single dorsiflexion without speed-accuracy goals or performance feedback. To sustain attention and motivation, all participants received standardized verbal encouragement (“do your best and strive for improvement in both speed and accuracy”) at the beginning of each rest period.

### Validation of the speed-accuracy function for motor skill

We assessed performance in the visuomotor task by utilizing the two-parameter speed-accuracy trade-off (SAF) model proposed by Reis et al. (2009). We included a dedicated speed-accuracy validation set to confirm that the present apparatus and task modifications preserved the expected lawful relationship between completion time (CT) and error rate (ER), thereby supporting construct validity and enabling a direct comparison with prior work using the same framework. Within this model, each trial produces an ER (the proportion of trials producing over or undershoot) at a designated CT. The relationship is characterized as follows:

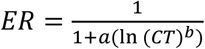

where *a* (the skill parameter) reflects the participants’ ability to execute fast, accurate movements, and *b* characterizes the slope of the tradeoff. CT denotes the prescribed movement time, and ER is the observed error rate (fraction of missed targets per block). To permit log-scale analyses, blocks with perfect accuracy (ER = 0) were assigned a nominal error rate of 0.01. To validate that the SAF model captured behavior in our visuomotor task and apparatus, we conducted a dedicated speed-accuracy protocol in an independent sample of 19 healthy adults (mean age 27.0 ± 3.1 years; 11 females). The participants completed nine pacing conditions, each consisting of one block of 10 trials, with the target movement time (MT) fixed within each block. The prescribed MT values were 2.4, 1.9, 1.3, 1.1, 0.9, 0.8, 0.75, 0.6, and 0.5 s, presented in randomized order across blocks. Consequently, each block lasted approximately 10 × MT seconds (i.e., ∼24 s at MT = 2.4 s to ∼5 s at MT = 0.5 s), excluding brief inter-trial transitions. MT was imposed by timing the appearance of successive targets at fixed intervals (one target every MT seconds). The participants were instructed to synchronize their movements to the target cadence (i.e., complete each movement within the prescribed MT) while still aiming to hit each target as accurately as possible. ER was computed per block as the proportion of trials that resulted in an over-or undershoot. Previous studies have used metronomic-paced cueing; however, because of the nature of visuomotor paradigms, we decided to pace the participants with quantifiable visual cuing, relying purely on visuo-motor interaction rather than including auditory sensory processing (Reis *et al*., 2009). Ten of these participants completed five days of practice, followed by an identical post-test. We fitted *a* and *b* to each individual’s baseline and post-test data using nonlinear least-squares regression (Fig. 1C). Examination of the resulting residuals showed no systematic deviation or trend, confirming an adequate model fit. Consistent with prior SVIPT findings, *b* remained stable (6.8% decrease), while *a* increased by 479% (Fig. 1D) with practice, validating the model for lower limb position control. For our main analyses, we isolated *a* using algebraic inversion:

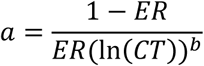

We then applied a natural logarithmic transform (log *a*, henceforth skill measure) to stabilize the variance. To characterize learning, we summed within-session skill gains over Days 1–5 (online learning) and overnight changes between sessions (offline learning). In all cases, *b* was fixed at 5.79, so that changes in skill reflect the modulation of the skill parameter *a* alone.

### Stimulation technique

tDCS was applied online during motor practice and delivered through two rubber electrodes, 3 mm thick; an anode (25 cm²) was positioned over the tibialis anterior hotspot identified via transcranial magnetic stimulation (TMS), oriented with the plugin posteriorly to maximize current flow along the central sulcus, and the cathode (35 cm²) was placed over the contralateral supraorbital region, Fp2/Fp1 according to the 10-20 EEG system. The electrodes were covered with 2 mm Ten20® conductive paste (Weaver and Company, Aurora, CO, USA). The paste was evenly distributed across the electrode surface using a specialized 3D-custom-made device. The electrodes were fixed on the head using headbands or rubber straps. Stimulation was delivered at 2 mA for 20 min per session with a 30 s ramp-up/down using a battery-driven, constant-current stimulator, Nurostym TES device (Brainbox, United Kingdom). Sham stimulation mimicked the initial ramp-up/down but did not deliver current throughout the session. The impedance measured directly between the stimulator and electrodes was kept below 5 kΩ. Prior to electrode placement, the skin surface was treated with Nuprep (Weaver and Company, Aurora, CO, USA), a mild abrasive gel that lowers skin impedance. The participants and the investigator performing the tests were blinded to the type of tDCS using a double-blind code for the tDCS stimulator. Note that the duration of the motor practice session varied according to the participants’ task performance (differences in completion time). Thus, stimulation was sometimes terminated before session completion but never after task completion. Non-overlap between session end and tDCS offset was 4.6 min in CON (fixed and paced by design), 4.27 ± 4.25 min in SKILL-STIM, and 4.80 ± 4.52 min in SKILL-SHAM (mean ± SD). Participants completed a questionnaire after the last training session to determine whether they thought they had received real or sham tDCS (Table 1). To estimate the E-field distribution induced by the montage, computational modeling was conducted using SimNIBS 4.0 (https://simnibs.github.io/simnibs) (SimNibs, DRCMR and DTU, Copenhagen and Lyngby, Denmark), deploying the ‘Ernie’ head model (Saturnino *et al*., 2015, 2019) (Fig. 1E). The magnitude of the electric field (normE) was evaluated within the gray matter (region index = 2). The peak E-field strength, defined as the 99.9th percentile, was 0.697 V/m. The 95th and 99th percentiles were 0.328 V/m and 0.43 V/m, respectively. Focality was assessed as the volume of gray matter (in mm³) exposed to at least 50% or 75% of the peak E-field intensity. This resulted in focality estimates of 27,000 mm³ and 3,290 mm³, respectively. These values indicate that while the field intensity peaks were localized, the stimulation also engaged relatively broad cortical areas, consistent with conventional tDCS focality profiles. Finally, region-of-interest (ROI) analysis at MNI coordinates (−4, 72, 46), corresponding to the lower limb representation in the M1 (Francis *et al*., 2009), revealed a mean electric field magnitude of 0.23 V/m, confirming that the leg motor area was effectively targeted.

### Electrophysiological recordings

#### Electromyography (EMG)

Surface electromyography (EMG) was recorded from the Tibialis Anterior (TA) and soleus (SOL) muscles of the non-dominant leg using Ag-AgCl electrodes (0.5 cm diameter, 1 mm between electrodes; Blue Sensor, Ambu Inc.,USA) in a bipolar montage across the muscle belly and a ground electrode mounted on the *patella*. Signals were amplified (×200), band-pass filtered (5 Hz–1 kHz), digitized at 2 kHz using a Cambridge Electronic Design system (CED 1401 Ltd., UK; Signal v6.05), and stored on a PC for off-line analysis. Line noise was suppressed using a 50 Hz Hum Bug eliminator (Digitimer).

#### Peripheral Nerve Stimulation

To elicit maximal compound muscle action potentials, the maximal M-wave or M_max_ in the TA muscle, electrical peripheral nerve stimulation was applied to the *common peroneal nerve* below the *caput fibulae* as a 200 µs pulse delivered by a constant-current stimulator (Digitimer DS27A, Digitimer, UK) through bipolar surface stimulation electrodes. Peripheral nerve stimulation was elicited at interstimulus intervals of 4 s, and the current was increased until the maximal peak-to-peak amplitude of the M-response was elicited. M_max_ was obtained immediately before generating TMS measurements, both before and after practice, to enable day-to-day comparisons (Day 1 and Day 5) and adjust for any practice-induced fatigue.

#### Transcranial Magnetic Stimulation

Single-pulse transcranial magnetic stimulations (TMS) were delivered to the contralateral hemisphere M1 of the performing leg via a figure-of-8 coil (MagPro MC-B70 Butterfly coil; MagVenture A/S, Denmark) connected to a MagPro X100 stimulator (MagVenture A/S, Denmark) for focused biphasic stimulation. To ensure that the orientation was optimal, the TA hotspot was identified through a mini-mapping procedure and defined as the location of the coil that elicited the largest and most consistent motor evoked potential amplitude (MEPs) during suprathreshold stimulations while the participant was at rest. During the mapping procedure, the coil was initially placed tangentially on the scalp at a 90° angle to the mid-sagittal plane, inducing medio-lateral to lateral-medio (ML-LM) current across the leg representation of the motor cortex. Subsequently, we rotated the coil to different orientations, from ML to posterior-anterior, which induced an antero-posterior followed by postero-anterior (AP-PA). ML coil orientation for the lower limb is suggested to induce a current perpendicular to the axonal direction of corticospinal neurons and the central sulcus, which is thought to be a relatively effective orientation for activating corticospinal output from the leg area by maximizing the induced electric field component normal to the cortical surface (Hand *et al*., 2020). The resting motor threshold (rMT) was defined as the stimulus intensity required to elicit recognizable MEPs with an amplitude above 0.05 mV in a minimum of five out of ten consecutive stimulations (Rothwell *et al*., 1999). The rMT assessed prior to motor practice on Day 1 was used exclusively to estimate the 120% rMT. Muscle relaxation was monitored using concurrent EMG recordings. Twenty TMS stimulations (120% of rMT) were delivered at each time point (prior to and after post-practice Day 1 and prior to and after post-practice Day 5) to measure MEP amplitudes. A neuronavigation system (Magstim TMS Patient Cap) was used to ensure the same positioning of the coil throughout the experiments. For a subset of participants (SKILL-STIM = 1, SKILL-SHAM = 1, CON = 4), rMT could not be reliably determined (MEPs ≥0.05 mV in 5/10) due to high rMT. In these cases, we identified the hotspot based on consistent responses of <0.05 mV and/or occasional suprathreshold MEPs ≥0.05 mV and proceeded with the same montage for tDCS placement. These participants also performed the behavioral/stimulation protocol described above.

### Data analysis and statistics

All statistical analyses were conducted using R (v4.1.3), primarily via the *brms* package (Bürkner, 2017), which interfaces with Stan (Carpenter *et al*., 2017) to implement Bayesian models using Hamiltonian Monte Carlo (HMC) sampling. Primary and exploratory outcome measures, skill measures, CT, and ER were analyzed. Skill learning dynamics were further decomposed into online learning (within-session improvements, e.g., last block – first block) and offline learning (between-session changes, e.g., first block of Day 2 - last block of Day 1). MEP amplitude was averaged across TMS stimulations for each participant. All signals were visually inspected during and after the experiment to exclude M-waves and MEPs exhibiting pre-stimulus EMG activity within the 100 ms preceding stimulation. Post hoc, outliers exceeding ±2 standard deviations from the mean were excluded. MEP amplitudes were normalized to the corresponding M_max_ to enable comparisons across sessions. Finally, as an exploratory analysis, we tested whether the baseline skill level moderated the effect of tDCS on the learning rate. Participants were divided into high and low baseline performers based on their initial baseline skill estimate (pre-practice). We then evaluated whether the estimated learning rate differed between active tDCS and sham within each subgroup, treating this analysis as hypothesis-generating, given reduced power and the post hoc subgrouping

#### Model specification

All models were fit as hierarchical Bayesian multilevel models, offering the flexibility of which different levels can have both constant and varying effects (fixed or random effects in frequentist statistics) and repeated measures nested into participants (Gelman & Hill, 2006). The ‘brm’ package uses formula syntax similar to that of the *lme4* package (Bates *et al*., 2015), allowing for the flexible specification of complex hierarchical models. Posterior distributions are estimated using Stan, which employs Markov Chain Monte Carlo (MCMC) sampling based on the HMC algorithm (Stan Development Team (2017c); Carpenter et al., 2017). For linear models, the outcome variable (e.g., skill measure) was regressed on group and time predictors, with a participant-specific intercept to account for repeated measures. Weakly informative Student-t priors centered at 0 were used for the intercept (α), slopes (β), and variance parameters, reflecting prior uncertainty while constraining the values within plausible ranges based on the outcome distribution. Models were fit using four chains with 20000 iterations each (2000 warmup), with adapt_delta = 0.99 and max_treedepth = 20, to ensure robust convergence. Model diagnostics and convergence were evaluated using the Gelman-Rubin statistic (*Ȓ*=1.00 for all parameters), and effective sample sizes (Bulk_ESS and Tail_ESS) indicated stable posterior estimates. Posterior predictive checks and trace plots were performed and confirmed that the model predictions were aligned with the observed data. To model learning trajectories over time, we fitted non-linear growth models. Several candidate functions were evaluated, including S-shaped, polynomial, and domain-specific exponential growth models (Bürkner, 2018; Williams *et al*., 2019). Model comparison was performed using LOO cross-validation. The growth model provided by Williams et al. (2018) revealed the best fit, which is why this model was chosen as a prediction model (See supplementary material for detailed information).

#### Bayesian inference

Bayesian inference was conducted using the ‘hypothesis’ function from the ‘brms’ package, which enables formal testing of directed and interval-based hypotheses on posterior distributions. For behavior, block-level skill measures were analyzed using hierarchical regression, with *group* as a predictor. To evaluate within-group changes in CSE, we estimated pre-to-post differences in MEP amplitudes normalized to M_max_ on Days 1 and 5 for each group. We pre-specified directional contrasts based on theory: SKILL-STIM > SKILL-SHAM (additive tDCS effect) and SKILL-SHAM > CON (benefit of skilled over non-skilled practice), with a primary focus on Post-practice and Follow-up. These were evaluated as one-sided Bayesian hypotheses for posterior contrasts, P(Δ>0). Baseline comparisons used two-sided tests (and ROPE for practical equivalence). Crucially, we kept the directional framing fixed regardless of raw trends to preserve confirmatory inferences and avoid post-hoc reversals. To interpret the practical relevance of the effects, we applied the Region of Practical Equivalence (ROPE) decision framework (Kruschke, 2018). This approach offers a more nuanced interpretation of posterior evidence and avoids the limitations of rigid, threshold-based rules (Etz *et al*., 2024). This is particularly valuable in studies involving subtle intervention effects or high between-subject variability, where practical significance must be contextualized within broader scientific and behavioral frameworks. ROPE region was based on the suggested standardization (Kruschke, 2018), and the determination often used, that is, 10% of the response scale. Posterior contrasts were calculated as the difference in posterior draws between groups or conditions and were transformed into standardized effect sizes (Cohen’s *d*) (Cohen, 1988; Kruschke, 2015; McElreath, 2020; Gelman *et al*., 2020). A ROPE of ± 0.1 was used for Cohen’s d, corresponding to a commonly accepted threshold for negligible effects and consistent with Kruschke’s recommendation for small effect size boundaries. Both the 95% highest density intervals (HDIs) and the percentage of the posterior distribution falling within the ROPE (ROPE%) were reported. Rather than applying a dichotomous decision rule (e.g., “accept” or “reject” equivalence), we interpreted ROPE% as a graded measure of evidence: 1) Posterior distributions centered near zero with a wide HDI and ROPE% > 5% suggest uncertainty but support practical equivalence. 2) Distributions centered away from zero with minimal posterior mass inside the ROPE (e.g., ROPE% < 5%) indicate strong evidence against practical equivalence.

## Results

Figure 2 summarizes the behavioral progression throughout the motor learning paradigm. Panel A illustrates changes in the primary skill measure across the five days of motor skill or non-skilled practice, combined with either real or sham tDCS, and during the seven-day follow-up retention test. Panels B-E display skill changes from baseline to post-practice and post-practice to follow-up. Panel F shows skill improvement in online learning (within-session) and offline learning (between-session consolidation). Panels G and H present the complementary performance metrics, CT and ER, captured at baseline, post-practice, and follow-up. In Figure 3, Panels A and B display the performance trajectories of high- and low-initial performers across the full training timeline and timepoints. The near-transfer and transfer effects to the untrained leg are shown in Panels A and B, respectively, in Figure 4. Together, these behavioral trajectories provide a foundation for subsequent Bayesian evaluation of group differences and learning dynamics. Bayesian evaluation of the skill measure, CT, and ER at different time points is presented in Table 2 and Supplementary Material S2.

**Figure 2:**
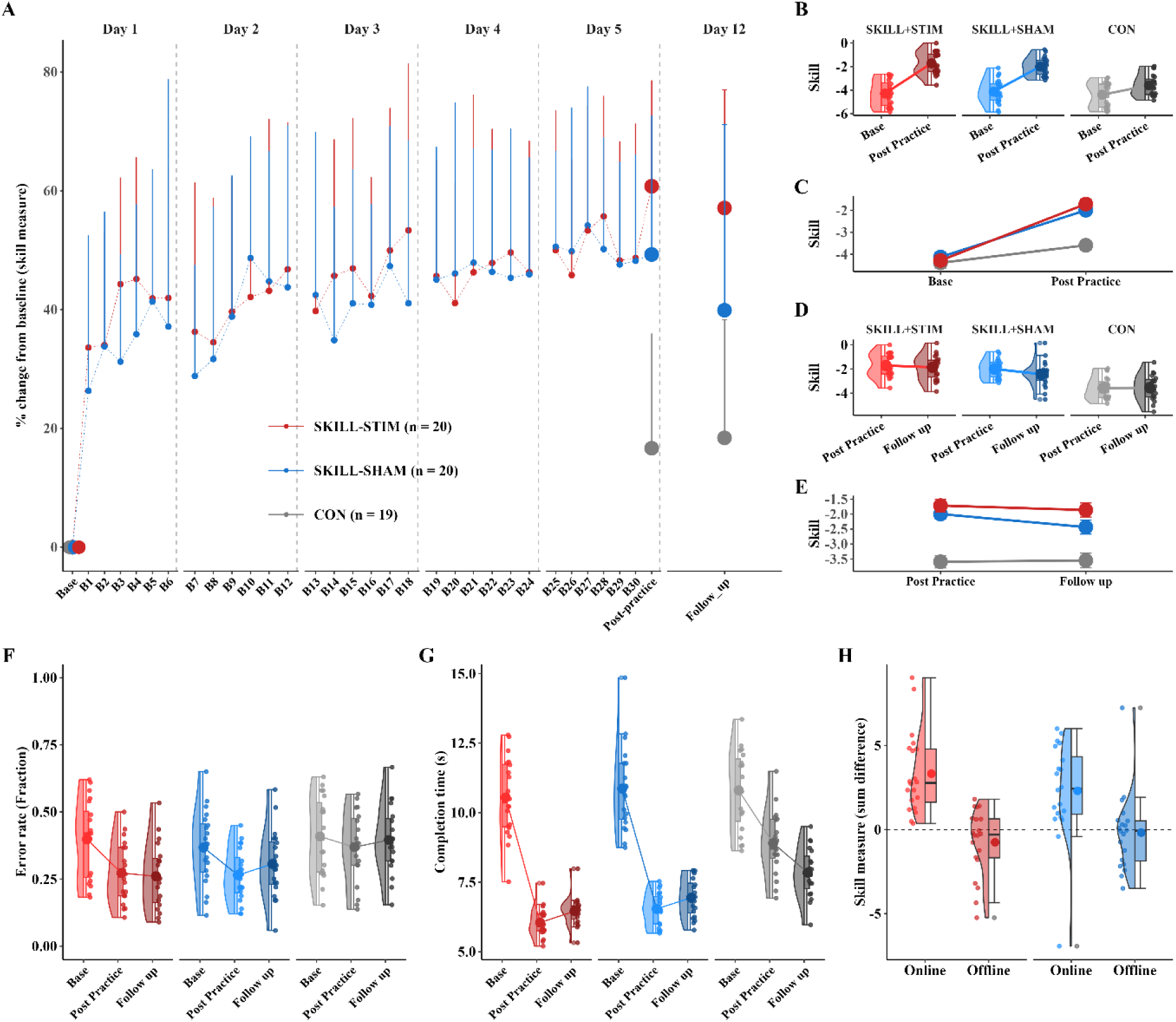
Behavior across practice and follow-up. A) Learning curves across blocks (% change from baseline). Groups: SKILL-STIM (red), SKILL-SHAM (blue), and CON (grey). Points show group means with error bars (SD); dashed lines connect blocks within day. B–E) Change in skill from Baseline to Post-practice and Post-practice to Follow-up by group (half-violin/box/points show distribution, mean, and spread). F-G) Component measures: Changes in Error Rate (ER) and Completion Time (CT) at Baseline, Post-practice, and Follow-up. H) Online (within-day) and offline (between-day) learning components for SKILL-STIM versus SKILL-SHAM. Colors: red = SKILL-STIM; blue = SKILL-SHAM; grey = CON. All panels use the same colors for the different groups.

**Figure 3:**
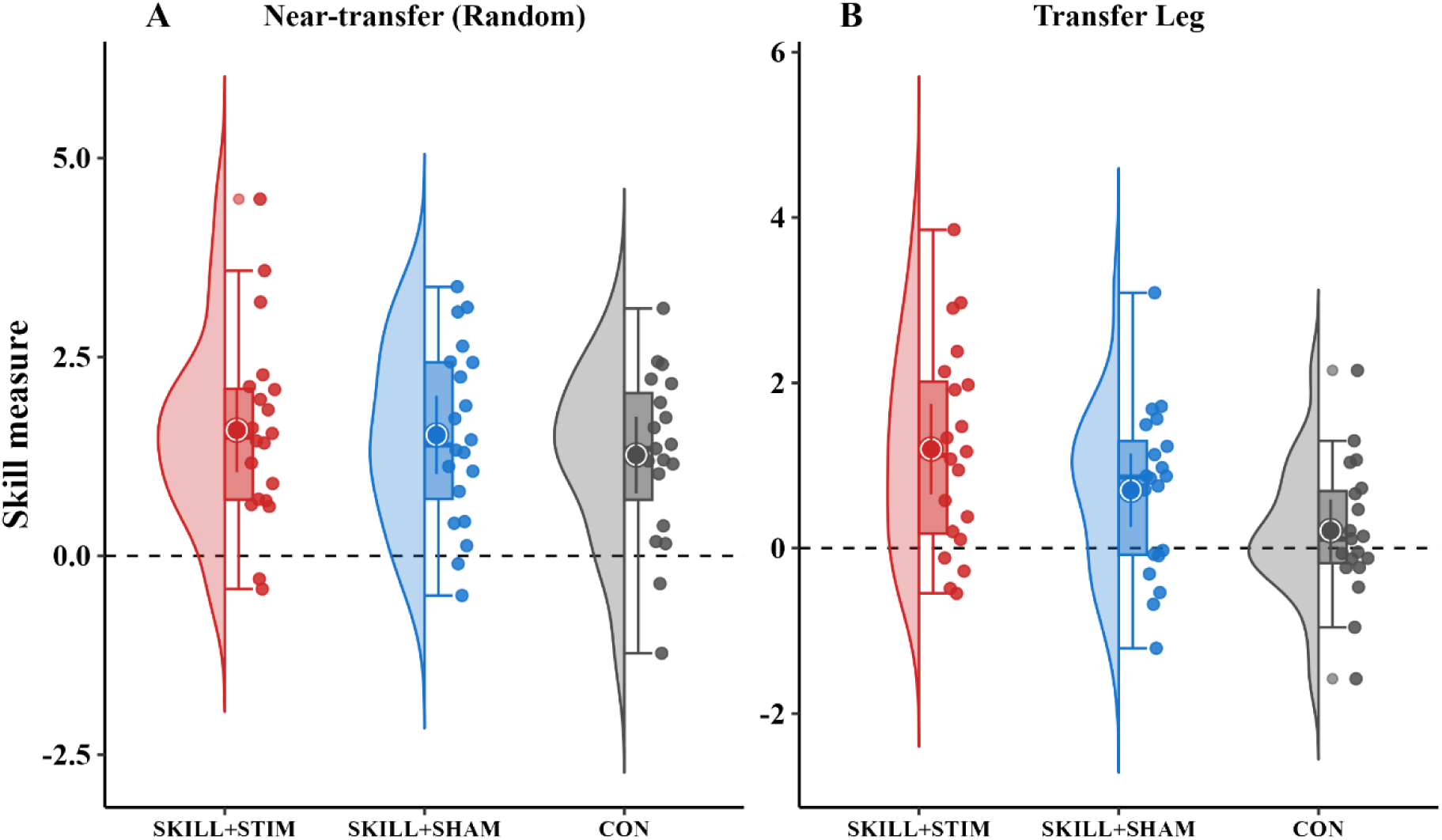
Transfer performance. Skill measures for the transfer effect in the random target transfer task (A) and the untrained leg (B). The y-axis is a skill measure log-transformed, which is used as a compound measure for speed and accuracy.

**Figure 4:**
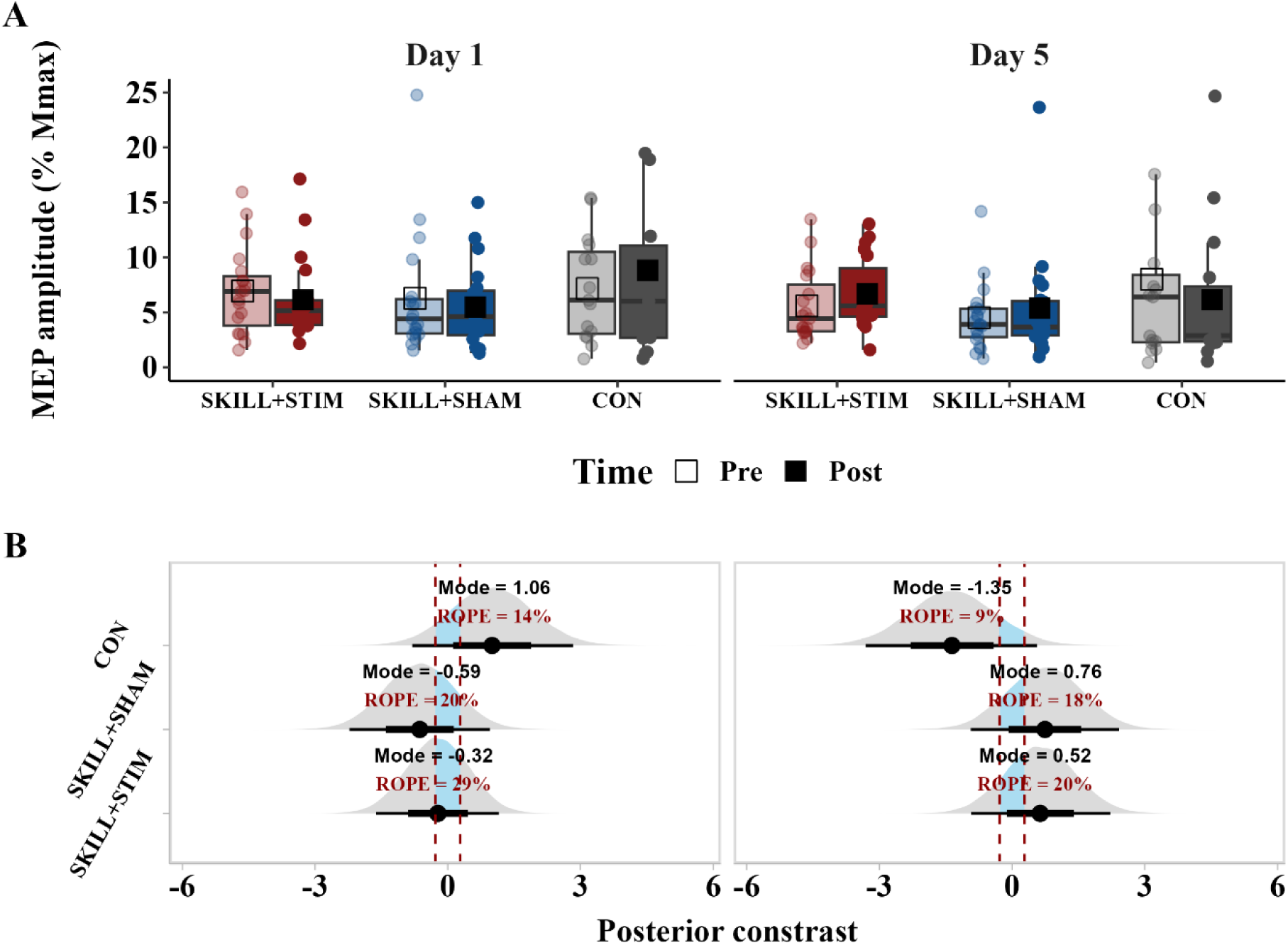
Corticospinal excitability. Boxplot distributions of MEP normalized to M_max_ measured before (pre) and after (post) a motor practice session on day one (left) and day five (right) for each group (A). Posterior contrasts of measures between pre vs. post for each group (B). The contrast reflects the difference in posterior predictions between Pre and Post at days one and five, with the 95% and 66% credible intervals as black bottom lines. The dashed vertical lines denote the Region of Practical Equivalence (ROPE) of ±10% of the standard deviation/sigma of the response variable, including the mode.

**Table 2:** Bayesian estimates for skill measure, completion time (CT), and error rate (ER). The estimate, standard error (SE), 95% high-density interval (95% HDI), and evidence ratio (Evid. Ratio), posterior probability, Region of Practical equivalence (ROPE), and Coheńs d effect size are presented in the table.

| Hypothesis testing | Measure | Estimate | SE | 95% HDI | Evid.Ratio | Posterior Probability | ROPE \|\| Cohen's *d* |
| --- | --- | --- | --- | --- | --- | --- | --- |
| No difference in baseline<br>(*SKILL-STIM* = *SKILL-SHAM*) | Skill | -0.13 | 0.27 | -0.63 \| 0.43 | 19.44 | 95% | 12% \| -0.3 |
|  | CT | -0.29 | 0.33 | -0.94 \| 0.34 | 9.18 | 90% | 8% \| -0.64 |
|  | ER | 0.03 | 0.04 | -0.05 \| 0.11 | 20.82 | 99% | 14% \| 0.35 |
| Differences between groups at post-practice<br>(*SKILL-STIM* > *SKILL-SHAM*) | Skill | 0.13 | 0.16 | -0.22 \| 0.42 | 3.08 | 75% | 17% \| 0.29 |
|  | CT | -0.22 | 0.28 | -0.67 \| 0.26 | 3.23 | 76% | 11% \| -0.49 |
|  | ER | -0.02 | 0.04 | -0.1 \| 0.04 | 2.68 | 73% | 15% \| -0.27 |
| Difference in nonlinearity/learning rate ($\gamma$)<br>(*SKILL-STIM* > *SKILL-SHAM*) | Skill | 0.01 | 0.39 | -0.63 \| 0.66 | 0.95 | 49% | 47% \| 0.01 |
|  | CT | -0.29 | 0.2 | -0.64 \| 0.05 | 12.03 | 92% | 6.6% \| -1.27 |
|  | ER | -0.23 | 0.82 | -1.6 \| 1.11 | 1.56 | 61% | 23% \| -0.01 |
| Differences between groups at follow-up<br>(*SKILL-STIM* > *SKILL-SHAM*) | Skill | 0.53 | 0.11 | 0.15 \| 0.91 | 106.78 | 99%* | 1% \| 1.13 |
|  | CT | -0.22 | 0.29 | -0.71 \| 0.24 | 3.67 | 79% | 10% \| -0.48 |
|  | ER | -0.08 | 0.04 | -0.15 \| 0 | 21.93 | 96%* | 4% \| -0.86 |
| No difference in baseline<br>(*SKILL-SHAM* = *CON*) | Skill | 0.21 | 0.2 | -0.2 \| 0.61 | 7.2 | 89% | 10% \| 0.48 |
|  | CT | -0.02 | 0.33 | -0.67 \| 0.61 | 8.46 | 89% | 12% \| -0.04 |
|  | ER | -0.04 | 0.04 | -0.13 \| 0.04 | 38.55 | 97% | 11% \| -0.44 |
| Differences between groups at post-practice<br>(*SKILL-SHAM* > *CON*) | Skill | 1.15 | 0.21 | 0.8 \| 1.48 | Inf | 100%* | 0% \| 2.6 |
|  | CT | -2.23 | 0.29 | -2.7 \| 1.76 | Inf | 100%* | 0% \| -4.92 |
|  | ER | -0.06 | 0.04 | -0.13 \| 0.01 | 10.93 | 92% | 7% \| -0.61 |
| Differences between groups at follow-up<br>(*SKILL-SHAM* > *CON*) | Skill | 0.59 | 0.21 | 0.24 \| 0.94 | 370 | 100%* | 0% \| 1.33 |
|  | CT | -0.79 | 0.29 | -1.24 \| -0.3 | 261.77 | 100%* | 0% \| -1.76 |
|  | ER | -0.05 | 0.04 | -0.12 \| 0.02 | 6.5 | 87% | 10% \| -0.52 |

### Skill measure

At baseline, the difference between SKILL-SHAM and CON showed support for practical equivalence, with 10% of the posterior inside the ROPE *(SKILL-SHAM = CON: 0.21 ± 0.20, 95% HDI [–0.20, 0.61], Evid. Ratio = 7.20, Post. Prob = 89%, ROPE = 10%, d = 0.48)*. After five days, the contrast increased substantially, with virtually no posterior mass within the ROPE and strong support for a group difference favoring the Sham group *(SKILL-SHAM > CON: 1.15 ± 0.21, 95% HDI [0.80, 1.48], Evid. Ratio = Inf ∞, Post. Prob = 100%, ROPE = 0%, d = 2.6)*. At follow-up, the difference remained in favor of SKILL-SHAM, with 0% of the posterior inside the ROPE *(SKILL-SHAM > CON: 0.59 ± 0.21, 95% HDI [0.24, 0.94], Evid. Ratio = 370, Post. Prob = 100%, ROPE = 0%, d = 1.33)* (Fig. 2A-E: Fig. 5A-B).

Comparing the SKILL-STIM and SKILL-SHAM groups, calculated as the contrast of posterior estimate, in skill measure was small and uncertain, with 12% of the posterior samples inside the ROPE range considered as evidence for practical equivalence *(SKILL-STIM = SKILL-SHAM: –0.13 ± 0.27, 95% HDI [–0.63, 0.43], Evid. Ratio = 19.44, Post. Prob = 95%, ROPE = 12%, d = −0.3)*. Following motor practice, the contrast favored the SKILL-STIM, but the evidence was inconclusive, with 17% of the posterior within the ROPE and a relatively wide credible interval, which provide evidence against a directional effect *(SKILL-STIM > SKILL-SHAM: 0.13 ± 0.16, 95% HDI [–0.22, 0.42], Evid. Ratio = 3.08, Post. Prob = 75%, ROPE = 17%, d = 0.29)*. No difference in learning rate was observed, which suggests no directionality *(SKILL-STIM > SKILL-SHAM: −0.01 ± 0.39, 95% HDI [–0.63, 0.66], Evid. Ratio = 0.95, Post. Prob = 49%, ROPE = 47%, d = −0.01)*. At follow-up, the posterior contrast favored the SKILL-STIM group with 1% of the distribution inside the ROPE, suggesting strong evidence for a meaningful group difference *(SKILL-STIM > SKILL-SHAM: 0.53 ± 0.11, 95% HDI [0.15, 0.91], Evid. Ratio = 106.75, Post. Prob = 99%, ROPE = 1%, d = 1.13)* (Fig. 2A-E: Fig. 5A-B).

For learning phase contrasts, online learning (i.e., within-session improvement) showed no meaningful difference between *SKILL-STIM and SKILL-SHAM*, with the posterior centered near zero and 20.4% of the distribution within the ROPE *(SKILL-STIM > SKILL-SHAM: –0.13 ± 0.58, 95% HDI [–1.08, 0.82], Evid. Ratio = 1.42, Post. Prob = 59%, ROPE = 20.4%)*. Similarly, offline learning (i.e., between-session consolidation) was practically equivalent across groups, with 15.7% of the posterior within the ROPE *(SKILL-STIM > SKILL-SHAM: 0.09 ± 0.78, 95% HDI [–1.17, 1.38], ER = 0.83, PP = 45%, ROPE = 15.67%)* (Fig. 2F).

Exploratory subgroup analyses suggested that baseline performance moderated the stimulation effect on the learning rate: active tDCS was associated with a steeper learning trajectory in high baseline performers but a shallower trajectory in low baseline performers (Fig. S3).

The comparison between SKILL-STIM and SKILL-SHAM in the near transfer random block indicated no evidence for a practical difference, with 15.8% of the posterior within the ROPE *(SKILL-STIM > SKILL-SHAM: 0.11 ± 0.21, 95% HDI [–0.24, 0.45], Evid. Ratio = 2.27, Post. Prob = 69%, ROPE = 15.8%, d = 0.23)*. Similarly, the contrast between SKILL-SHAM and CON was inconclusive, though modestly shifted toward Sham *(SKILL-SHAM > CON: 0.24 ± 0.21, 95% HDI [–0.10, 0.59], Evid. Ratio = 6.97, Post. Prob = 87%, ROPE = 8.95%, d = 0.5)* (Fig. 3A).

In the transfer leg task (baseline vs. post-practice), the contrast between *SKILL-STIM and SKILL-SHAM* suggested practical equivalence, with 10.2% of the posterior within the ROPE *(SKILL-STIM > SKILL-SHAM: 0.22 ± 0.21, 95% HDI [–0.12, 0.56], Evid. Ratio = 5.58, Post. Prob = 85%, ROPE = 10.18%, d = 0.49)*. The contrast between SKILL-SHAM and CON, by contrast, strongly supported a directional effect favoring Sham, with 0% of the posterior within the ROPE *(SKILL-SHAM > CON: 0.49 ± 0.21, 95% HDI [0.14, 0.83], Evid. Ratio = 92.26, Post. Prob = 99%, ROPE = 0%, d = 2.7)* (Fig. 3B).

### Completion Time

At baseline, the contrast in CT between the *SKILL-STIM and SKILL-SHAM* groups was small and uncertain, with 8% of the posterior distribution falling within the ROPE, suggesting strong evidence against practical equivalence *(SKILL-STIM = SKILL-SHAM: –0.29 ± 0.33, 95% HDI [–0.94, 0.34], Evid. Ratio = 9.18, Post. Prob = 90%, ROPE = 8%, d = –0.64)*. After motor practice, the contrast did not change accordingly, with 11% of the posterior inside the ROPE, again suggesting no practical difference *(SKILL-STIM < SKILL-SHAM: –0.22 ± 0.28, 95% HDI [–0.67, 0.26], Evid. Ratio = 3.23, Post. Prob = 76%, ROPE = 11%, d = –0.49)*. The follow-up comparison remained consistent, indicating a persistent contrast *(SKILL-STIM < SKILL-SHAM: –0.22 ± 0.29, 95% HDI [–0.71, 0.24], Evid. Ratio = 3.67, Post. Prob = 79%, ROPE = 10%, d = –0.49)*. For the learning rate, the contrast between groups was also in favor of the SKILL-STIM group, with a limited posterior mass in the ROPE *(SKILL-STIM < SKILL-SHAM: –0.29 ± 0.20, 95% HDI [–0.64, 0.05], Evid. Ratio = 12.03, Post. Prob = 92%, ROPE = 6.6%, d = –1.27.)* (Fig. 2G: 5C-D).

The baseline contrast in completion time between the SKILL-SHAM and CON groups was negligible, with a great amount of the posterior distribution falling inside the ROPE *(SKILL-SHAM = CON: – 0.02 ± 0.33, 95% HDI [–0.67, 0.61], Evid. Ratio = 8.46, Post. Prob = 89%, ROPE = 12%, d = –0.04)*. At post-practice, SKILL-SHAM completed the task substantially faster than the CON group, with 0% of the posterior inside the ROPE and clear evidence of a group difference *(SKILL-SHAM < CON: – 2.23 ± 0.29, 95% HDI [–2.70, –1.76], Evid. Ratio = >100, Post. Prob = 100%, ROPE = 0%, d = – 4.92)*. This contrast remained large at follow-up *(SKILL-SHAM < CON: –0.79 ± 0.29, 95% HDI [–1.24, –0.33], Evid. Ratio = 261.77, Post. Prob = 100%, ROPE = 0%, d = –1.76)* (Fig. 2G: 5C-D).

### Error Rate

At baseline, the contrast in ER between *SKILL-STIM and SKILL-SHAM* was close to zero, with 14% of the posterior falling within the ROPE, indicating modest evidence for practical equivalence *(SKILL-STIM = SKILL-SHAM: 0.03 ± 0.04, 95% HDI [–0.05, 0.11], Evid. Ratio = 20.82, Post. Prob = 99%, ROPE = 14%, d = 0.35)*. After practice, the contrast slightly favored the SKILL-STIM group, but the distribution remained uncertain and largely within the ROPE *(SKILL-STIM < SKILL-SHAM: – 0.02 ± 0.04, 95% HDI [–0.10, 0.04], Evid. Ratio = 2.68, Post. Prob = 73%, ROPE = 15%, d = –0.27)*. At follow-up, the contrast was stronger, with only 4% of the posterior inside the ROPE and high posterior probability, supporting a small but meaningful difference *(SKILL-STIM < SKILL-SHAM: – 0.08 ± 0.04, 95% HDI [–0.15, 0.00], Evid. Ratio = 21.93, Post. Prob = 96%, ROPE = 4%, d = –0.86)*. For the learning rate, the group difference was highly uncertain and largely inside the ROPE *(SKILL-STIM < SKILL-SHAM: –0.23 ± 0.82, 95% HDI [–1.60, 1.11], Evid. Ratio = 1.56, Post. Prob = 61%, ROPE = 23%, d = –0.01)* (Fig. 2H: 5E-F).

Between the *SKILL-SHAM and CON* groups, the baseline contrast was small and mostly outside the ROPE *(SKILL-SHAM = CON: –0.04 ± 0.04, 95% HDI [− 0.13, 0.04], Evid. Ratio = 38.55, Post. Prob = 97%, ROPE = 11%, d = –0.44)*. At post-practice, the contrast showed lower error rates for SKILL-SHAM, with a majority of the posterior outside the ROPE *(SKILL-SHAM < CON: –0.06 ± 0.04, 95% HDI [–0.13, 0.01], Evid. Ratio = 10.93, Post. Prob = 92%, ROPE = 7%, d = –0.61)*. This difference was maintained at follow-up *(SKILL-SHAM < CON: –0.05 ± 0.04, 95% HDI [–0.12, 0.02], Evid. Ratio = 6.5, Post. Prob = 87%, ROPE = 10%, d = –0.52)* (Fig. 2H: 5E-F).

### Corticospinal excitability

For the MEP amplitudes, hypothesis testing and ROPE-based equivalence testing were conducted to assess within-group pre–post changes on Days 1 and 5. At Day 1, the contrast in MEP amplitudes from pre to post in SKILL-STIM showed a mode of −0.62 with 11% posterior mass inside the ROPE, suggesting no change in group-level MEP amplitude *(SKILL-STIM pre–post: −0.32± 0.71, 95% HDI [–1.39, 0.94], Evid. Ratio = 0.6, Post. Prob = 38%, ROPE = 29%, d = −0.11)*. In the SKILL-SHAM group, the distribution was highly uncertain and largely overlapping zero, with more posterior mass inside the ROPE *(SKILL-SHAM pre–post: −0.59± 0.81, 95% HDI [–1.96, 0.7], Evid. Ratio = 0.28, Post. Prob = 22%, ROPE = 20%, d = −0.21)*. The CON group showed a nearly null effect *(CON pre–post: - 1.06± 0.93, 95% HDI [–0.51, 2.54], Evid. Ratio = 6.25, Post. Prob = 86%, ROPE = 14%, d = 0.38)* (Fig. 4A-B left column).

At Day 5, SKILL-STIM exhibited a posterior distribution shifted clearly above zero with minimal overlap with the ROPE, providing evidence for an increase in corticospinal excitability but with high uncertainty *(SKILL-STIM pre–post: 0.52± 0.8, 95% HDI [–0.68, 1.97], Evid. Ratio = 3.72, Post. Prob = 79%, ROPE = 20%, d = 0.19)*. The SKILL-SHAM group showed a smaller and more uncertain change, with moderate posterior mass inside the ROPE *(SKILL-SHAM pre–post: 0.76± 0.86, 95% HDI [–0.67, 2.15], Evid. Ratio = 4.16, Post. Prob = 81%, ROPE = 18%, d = 0.27)*. For the CON group, the distribution remained centered near zero *(CON pre–post: −1.35± 0.99, 95% HDI [–2.99, 0.26], Evid. Ratio = 0.09, Post. Prob = 8%, ROPE = 9%, d = −0.48)* (Fig. 4A-B right column).

The same analysis was conducted for changes in M_max_ from pre to post on Days 1 and 5. For the SKILL-STIM group on Day 1, the estimated change in M_max_ from pre to post was small and uncertain *(SKILL-STIM pre–post: −0.19 ± 0.18, 95% HDI [–0.56, 0.17], Evid. Ratio = 13.79, Post. Prob = 93%, ROPE = 14.03%)*. Approximately 14% of the posterior distribution fell within the ROPE, indicating weak evidence against practical equivalence. The SKILL-SHAM group showed a slightly larger decrease but with similar uncertainty *(SKILL-SHAM pre–post: −0.33 ± 0.20, 95% HDI [–0.71, 0.05], Evid. Ratio = 5.48, Post. Prob = 85%, ROPE = 5.65%)*. For the CON group, the posterior difference was likewise uncertain, with substantial overlap with zero *(CON pre–post: −0.20 ± 0.21, 95% HDI [–0.61, 0.2], Evid. Ratio = 7.89, Post. Prob = 89%, ROPE = 12.98%)*.

At Day 5, the SKILL-STIM group again showed a small positive shift, with the posterior mostly outside the ROPE, indicating no evidence of a meaningful decrease *(SKILL-STIM pre–post: −0.26 ± 0.19, 95% HDI [–0.64, 0.11], Evid. Ratio = 13.95, Post. Prob = 93%, ROPE = 9.12%)*. The SKILL-SHAM group showed a comparable effect *(SKILL-SHAM pre–post: −0.20 ± 0.19, 95% HDI [–0.58, 0.17], Evid. Ratio = 19.61, Post. Prob = 95%, ROPE = 13.18%)*. The CON group showed slightly more uncertainty but a similar trend *(CON pre–post: −0.32 ± 0.21, 95% HDI [–0.74, 0.1], Evid. Ratio = 6.47, Post. Prob = 87%, ROPE = 6.74%)*.

## Discussion

In this study, we compared five days of lower-limb motor skill practice with non-skilled ankle movements and investigated whether pairing practice with active tDCS adds benefit to motor learning. Practice alone produced clear gains in the skill measure relative to control movements and transferred to the untrained leg but not to an untrained sequence. Adding tDCS improved long-term retention without affecting online (within-session) or offline (between-session) performance changes across training days. However, exploration of the moderating effects of baseline performance revealed that tDCS increased learning rates in individuals with good baseline performance and decreased learning rates in individuals with low baseline performance. There was no consistent pattern in how the amplitude of MEPs obtained at rest changed within or between days, and the changes were not associated with changes in motor performance. Taken together, these findings indicate that while multi-session tDCS did not accelerate acquisition, it may have stabilized learning over time.

### Multiple days of lower limb practice combined with tDCS improved long-term retention of skilled sequence learning

Lasting improvements in visuomotor skill performance following motor practice (i.e., retention) require structural changes in the neural circuitry involved in executing the skill. Long-term retention of practiced motor skills is crucial for maintaining high-level performance and function. We used a modified version of the SVIPT task to investigate the effect of sequential skill practice against non-skilled control movements and the potential additive effects of tDCS on sequential skill acquisition and long-term retention. We show that multi-day practice paired with tDCS yields superior one-week retention of lower-limb skill learning, that is, less behavioral decay from post-practice to follow-up. This extends prior work on upper-limb tasks by demonstrating a retention benefit for skill learning involving the leg and helps reconcile mixed evidence on skill retention. Several studies have reported enhanced long-term retention with anodal tDCS (Saucedo Marquez *et al*., 2013; Pixa *et al*., 2017; Talimkhani *et al*., 2018), whereas others have found no effect (Ciechanski *et al*., 2017). Work using a related multiday paradigm also revealed positive retention effects at five different time points from the end of motor practice to three months after, but this effect was evident already during the five days of motor practice and did not occur as a specific enhancement of long-term retention (Reis *et al*., 2009). Notably, the retention advantage observed in the present study emerged without reliable changes in resting MEPs, either acutely or across days. This dissociation suggests that tDCS can bias consolidation processes without necessarily shifting corticospinal excitability. A plausible account is that weak, behavior-contingent fields modulate late-phase plasticity (e.g., protein-synthesis–dependent LTP) that unfolds over hours to days (Brashers-Krug *et al*., 1996; Rioult-Pedotti *et al*., 2000; Robertson, 2005; Luft & Buitrago, 2005; Rosenkranz *et al*., 2007; Kantak & Winstein, 2012), and/or influence sequence-specific representations in cortico-striatal loops that are not optimally captured by single-pulse resting TMS (Debas *et al*., 2014). Collectively, these findings indicate that pairing practice with tDCS can enhance the durability of lower-limb motor skill learning, a target with clear translational relevance for athletic training and rehabilitation. The absence of reliable changes in resting MEPs suggests that these retention effects are not necessarily expressed as a sustained increase in corticospinal excitability at rest.

### tDCS across multiple days did not boost learning during or between practice days

Contrary to our hypothesis and previous findings (Reis et al., 2009), five consecutive days of anodal tDCS targeting M1-Leg delivered during motor practice did not enhance online or offline learning compared to sham stimulation. This finding diverges from seminal SVIPT studies in the upper limb, which reported enhanced learning rates with concurrent tDCS, particularly in the cumulative offline learning phase between practice sessions (Reis et al., 2009; Reis & Fritsch, 2011). However, the present study differs from previous studies in several important methodological aspects. First, we employed a dynamic, sequential position-controlled motor task of the lower limb, manipulating both the effector and task control parameters. Most prior studies have focused on facilitating learning during upper-limb tasks involving finger tapping and precision grips (Nitsche *et al*., 2003; Fan *et al*., 2017; Greeley *et al*., 2020; Kunaratnam *et al*., 2022). For an overview, see Nielsen et al. (2025), including work using a comparable sequential precision task (Reis et al., 2009). We would expect lower limb practice to benefit from concurrent tDCS comparably, as precise control of dorsiflexion is similar to that of the hand, contingent on corticospinal and corticomotoneuronal pathways (Brouwer & Ashby, 1992; Nielsen *et al*., 1993; Perez *et al*., 2006). However, relatively few studies have targeted M1-Leg during ankle visuomotor training, and those that have reported no selective offline gains (Sriraman *et al*., 2014; Foerster *et al*., 2018). Moreover, position- versus force-control tasks differentially engage corticospinal circuits (Nielsen *et al*., 2025b), which may further modulate responsiveness to tDCS; however, Foerster et al. (2018) also found null effects in a force-control paradigm. An interesting exploratory finding was reported by Foerster et al. (2018). Here, consolidation was enhanced only for participants in whom an MEP could be evoked from the TA during rest, which relates to the scalp-to-cortex distance as well as the mesial depth of the M1-Leg representation. Although we cannot reject the possibility that trained effectors or the demand for position rather than force control resulted in the absence of offline effects between practice days, we would expect that sufficiently strong tDCS of M1-Leg should impact the corticospinal control of dorsiflexion similarly to the way anodal stimulation of M1-Hand would interact with practice-dependent plasticity during and following hand practice.

In the present study, we employed a dynamic lower-limb version of the SVIPT task. In contrast to previous studies, we evaluated whether learning was specific to the trained sequence and limb. Interestingly, we found that performance increments in an untrained sequence were comparable across all three groups. We conclude that skilled sequence practice did not improve generalizable visuomotor control more than what can be expected from test-retest effects, and that concurrent tDCS did affect visuomotor control. The sequential nature of learning is further supported by the finding that sequence-specific improvement with skilled practice transferred to the untrained leg. Collectively, these two findings suggest that the practice introduced a skilled sequence representation accessible for employment by both legs. Although the absence of transfer (sequence and interlimb) tests in the existing literature impedes a direct comparison, it is noteworthy that Reis et al. (2009) reported clear effects on offline learning when training on a single sequence. In contrast, Forster et al. (2018) and Sriraman et al. (2014) both used force and position targets in a random order. Consequently, we consider it unlikely that the sequential nature of the task impeded the between-session improvement.

Finally, as highlighted by Forster et al. (2018), it is challenging to direct a sufficiently strong E-field to the M1-LEG selectively. Given the position of M1-Leg deep in the mesial wall, it is worth questioning whether the intensity of the induced e-field was of sufficient strength at the target ROI. In the exemplary e-field computation (Fig. 1E), we found a peak strength of 0.697 V/m, which is within the lower range of intensities that can bias neural activity. Weak DCS in vitro field intensities have been shown to alter synaptic plasticity and can be a functional targeting tool to enhance ongoing neural activity (Kronberg *et al*., 2020; Sharma *et al*., 2022). In line with the well-known lack of focality from conventional tDCS, and as evident from Figure 1E, e-field intensity was most likely as high or higher in other cortical areas. In support, our exemplary simulation revealed that the e-field intensity in medial area 6, an area typically implicated in sequence learning, as defined by coordinates taken from Hardwick et al. (2013) and Sami et al. (2014), was stimulated with 0.3 V/m. Although a discussion of the potential interaction between motor practice and co-stimulation of all cortical and subcortical areas receiving electrical stimulation above the threshold for neuromodulation is beyond the scope of this study, caution is warranted in interpreting our null findings, as the theory could arise from opposing effects from co-stimulation of non-targeted networks.

An interesting result from the exploratory analysis was that high-baseline performing participants benefited more from active tDCS than from sham tDCS, exhibiting steeper learning trajectories. In contrast, tDCS did not benefit learning rates for participants with the lowest baseline performance. This unexpected result challenges the assumption that low performers, who arguably have more room for improvement, are more responsive to neuromodulation (Sánchez-Kuhn *et al*., 2018; Vergallito *et al*., 2022). Instead, our findings suggest that baseline performance level may interact with tDCS in complex and non-linear ways, warranting further investigation of individual variability in stimulation responsiveness.

### No reliable CSE modulation, consistent with sequence-specific learning outside M1

Across groups, no practice-dependent modulation of resting MEPs was observed after practice on day one or day five. Considering behavior and improvement on the trained sequence without advantage on the near-transfer (random) block, these findings favor a sequence-specific account of learning rather than a generalized increase in motor acuity, where demands for speed and accuracy have been demonstrated to increase CSE (Christiansen et al., 2018, 2021). Converging findings from animal studies indicate that once a sequence is well learned, its execution can be supported predominantly by subcortical circuits, the sensorimotor striatum, and thalamo-striatal pathways, while M1 becomes less necessary for expression (Kawai *et al*., 2015; Dhawale *et al*., 2021; Wolff *et al*., 2022; Mizes *et al*., 2024). In such a regime, resting M1-based MEPs provide a relatively insensitive window into the locus of plastic change, which helps reconcile robust sequence-bound behavioral gains with the absence of group-level CSE effects.

This interpretation also places our findings within the context of previous CSE results for lower-limb skill practice. A seminal study reported that a single session of position-control practice of randomized foot flexion series increased CSE when targeting the M1 leg area and improved accuracy relative to non-task-specific movement (Perez *et al*., 2004), which may be a result of an increase in motor acuity. As emphasized by a recent lower-limb review (Woodhead *et al*., 2024), leg-area recordings face additional variability from coil-target geometry and depth, and resting single-pulse MEPs may be less sensitive than input–output curves, paired-pulse metrics, or task-engaged/active-contraction probes as compared to effects in the M1 hand knob area due to neuroanatomical and morphometric reasons. More broadly, resting MEPs are influenced by arousal, attention, and expectancy (Bestmann & Krakauer, 2015), which modulate corticomotor communication without necessarily indexing durable synaptic changes.

Although anodal tDCS applied to M1 is often associated with acute increases in CSE after a single session (Nitsche & Paulus, 2001; Ambrus *et al*., 2016) but also with varying results (Chew *et al*., 2015; Madhavan *et al*., 2016), our multi-day protocol showed no reliable resting MEP modulation on Day 1 or Day 5. Recent work illustrates increased MEP amplitudes across multi-session finger-sequence learning, regardless of stimulation intensity (0, 4, and 6 mA) and without a clear link to performance (Hsu *et al*., 2025). Their design lacked transfer control; therefore, broader motor acuity versus sequence-specific gains could not be disambiguated. At the mechanistic level, in-vitro work using lower, human-relevant electric fields still demonstrates synaptic plasticity (Sharma *et al*., 2022), supporting the plausibility that weak fields can bias active circuits even if resting MEPs remain unchanged. Taken together, the absence of practically meaningful resting-MEP change, coupled with sequence-specific behavioral improvement, is consistent with plasticity consolidating outside M1 and/or manifesting in task-dependent neural states not captured by resting TMS.

## Conclusion

We demonstrate that motor skill practice, relative to non-skilled control movements, produced robust learning, interlimb transfer, and superior performance at follow-up. Adding active tDCS during practice did not amplify online (within-session) or offline (between-session) gains during training but selectively strengthened the long-term retention of a complex discrete motor task requiring precise and rapid ankle movements under high visuomotor demands. Baseline performance level emerged as an important factor in determining the extent to which tDCS could augment skill acquisition, with higher-performing individuals deriving the greatest benefit from concurrent stimulation and practice. Together, these findings indicate that concurrent tDCS may preferentially support skill retention rather than immediate performance, whereas practice itself drives the bulk of acquisition and transfer processes. Practically, this supports the use of transcranial direct current stimulation as an adjunct to promote retention of learned lower-limb skills, helping inform timing and dosing in rehabilitation and performance settings while emphasizing that well-designed practice remains the primary driver of improvement.

## DATA AVAILABILITY

The source data that support the findings of this study are available from the corresponding author upon reasonable request.

## SUPPLEMENTAL MATERIAL

Supplementary material – S1: Convergence assessment for Bayesian models.

## AUTHOR CONTRIBUTIONS

A.L.K., A.N.K., L.C., and J.L.-J. conceived and designed the study; A.L.K. performed the experiments; A.L.K. analyzed the data; A.L.K., A.N.K., L.C., and J.L.-J. interpreted the results of the experiments; A.L.K. prepared the figures; A.L.K. drafted the manuscript; A.L.K., A.N.K., L.C., L.J., J.R.B., and J.L.-J. edited and revised the manuscript: A.L.K., A.N.K., L.C., L.J., J.R.B., and J.L.-J. approved the final version of the manuscript.

## ACKNOWLEDGMENTS

We thank all the participants and acknowledge Team Danmark and the Novo Nordisk Foundation for funding this study.

