## Supplementary material for "Transcranial Direct Current Stimulation enhances long-term retention after 5 days of lower-limb motor skill learning"

### Supplementary S1 – Statistical details

**Model specification**

All models were fit as hierarchical Bayesian multilevel models, offering the flexibility of which different levels can have both constant and varying effects (fixed or random effects in frequentist statistics) and repeated measures nested into participants (Gelman & Hill, 2006). A key strength of Bayesian statistics is the explicit representation of uncertainty; model parameters are treated as probability distributions rather than fixed values. Inference proceeds by updating prior beliefs (priors) in light of the observed data (likelihood), yielding a posterior distribution that reflects both sources of information (Kruschke, 2015). Even with less informative priors, Bayesian modeling has several advantages, including the intuitive nature of Bayesian inference as a probability function referring to the experience of uncertainty rather than a limit of relative frequency (Nalborczyk et al., 2019). The ´brms´ package uses formula syntax similar to that of the *lme4* package (Bates et al., 2015), allowing for the flexible specification of complex hierarchical models. Posterior distributions are estimated using Stan, which employs Markov Chain Monte Carlo (MCMC) sampling based on the HMC algorithm. This method enables efficient sampling from high-dimensional parameter spaces and tends to converge more quickly and reliably than traditional MCMC, particularly when the model parameters are strongly correlated (Stan Development Team (2017c); Carpenter et al., 2017). For linear models, the outcome variable (e.g., skill measure) was regressed on group and time predictors, with a participant-specific intercept to account for repeated measures. The basic model structure can be written as:

*y_i_ ∼ Student-t(μ_i_,σ,ν)*

*μ_i_ =α + α_participant[i]_ + β_1_⋅ Group_i_ + β_2_⋅ Time_i_ + β_3_⋅ (Group_i_ × Time_i_)*

where *α_participant_ ∼ N*(0, *σ_participant_*), and weakly informative priors were used.

*α ∼ Student-t(3, -3.4, 2.5)*

*β_k_ ∼ Student-t(3, 0, 2.5)*

*σ, σ_participant_ ∼ Student-t(3, 0, 2.5)*

The first two lines define the likelihood and linear model. The subsequent lines specify the priors for the model parameters. Weakly informative Student-t priors centered at 0 were used for the intercept (α), slopes (β), and variance parameters, reflecting prior uncertainty while constraining the values within plausible ranges based on the outcome distribution. Models were fit using four chains with 20000 iterations each (2000 warmup), with adapt_delta = 0.99 and max_treedepth = 20, to ensure robust convergence. The Student-t family was specified to allow heavier tails and accommodate potential outliers. Model diagnostics and convergence were evaluated using the Gelman-Rubin statistic (*Ȓ*=1.00 for all parameters), and effective sample sizes (Bulk_ESS and Tail_ESS) indicated stable posterior estimates. Posterior predictive checks and trace plots were performed and confirmed that the model predictions were aligned with the observed data. To model learning trajectories over time, we fitted non-linear growth models. Several candidate functions were evaluated, including S-shaped, polynomial, and domain-specific exponential growth models (Bürkner, 2018; Williams et al., 2019). Model comparison was performed using LOO cross-validation. The growth model provided by Williams et al. (2018) revealed the best fit, which is why this model was chosen as a prediction model and can be expressed as:

*μ_i_ = β + (α − β) ⋅ exp(− exp(γ)⋅ time_i_)*

Where:

α = initial performance (baseline skill), β = final performance (asymptote), and γ = learning rate. Each of these parameters was modeled hierarchically across subjects and as a function of group:

*α ∼ Group+(1∣Participant)*

*β ∼ Group+(1∣Participant)*

*γ ∼ Group+(1∣Participant)*

The priors were weakly informative and grounded in the previous literature.

*α ∼ N(−4.5, 1), β ∼ N(−2, 1), γ ∼ N(−1.5, 1)*

*Group effects ∼ N(0, 1)*

*Random effects ∼ Student-t(3, 0, 2)*

**Directionality in hypothesis testing**

We pre-specified directional contrasts based on theory and prior work. Our primary tests asked whether SKILL-STIM > SKILL-SHAM (additive effect of tDCS during practice) and SKILL-SHAM > CON (benefit of skilled practice over non-skilled movements), with particular focus on Post-practice and Follow-up. These were evaluated as one-sided Bayesian hypotheses on the posterior contrasts (e.g., P[Δ>0]). For baseline comparisons, where no a priori group differences were expected, we used two-sided tests (and, when relevant, an interval/ROPE to assess practical equivalence). Importantly, we kept the directional framing fixed regardless of the visual trend in raw means: even when a posterior mode was near zero or reversed, we interpreted results within the predefined directional hypotheses. This preserves confirmatory inference, avoids post-hoc reversals, and aligns with best practices for Bayesian confirmatory analysis.

### Supplementary S2 – Results and posterior predictive

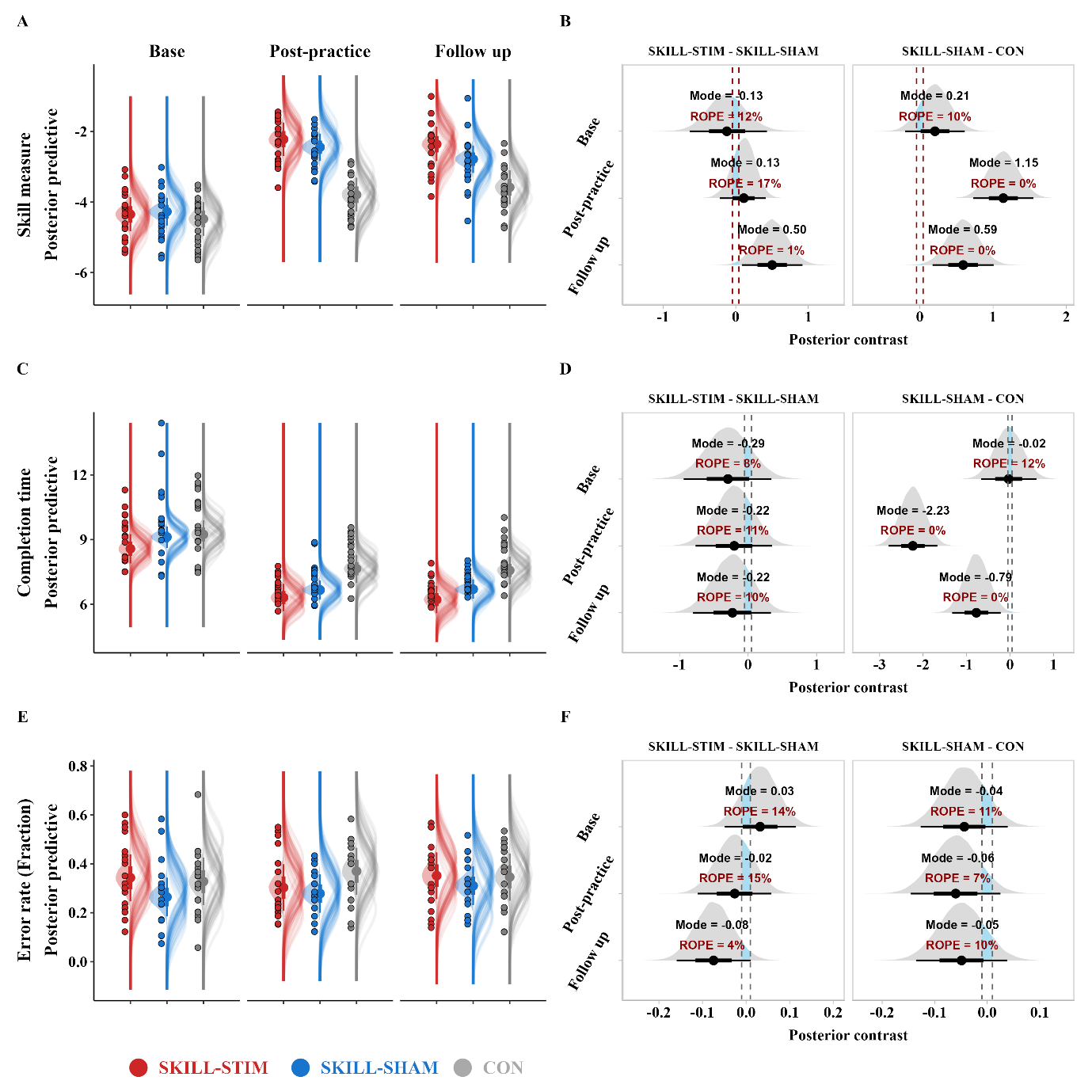

***Figure S2: Posterior predictive.*** *Skill measure for each group across three key timepoints: Baseline (Base), Post-practice, and Follow-up (A, B), and for completion time and error rate (C-F). (A, C, E) Half-eye plots represent the model-based estimated posterior distribution (. epred) for each group at each time point. The outer shaded area indicates the 95% credible interval, and the inner density indicates the 66% credible interval. The overlaid dots represent the observed values of the individual parameters. The outlines (gray ridges) visualize the 40 samples from the predicted posterior distributions. (B, D, F) Posterior contrasts of skill measure between (1) SKILL-STIM vs SKILL-SHAM and (2) SKILL-SHAM vs CON groups at different time points. The contrast reflects the difference in posterior predictions between groups. Half-eye plots show the posterior distributions of these differences, with 95% and 66% credible intervals shaded. The dashed vertical lines denote the Region of Practical Equivalence (ROPE).*

### Supplementary S3 – High vs. low performers

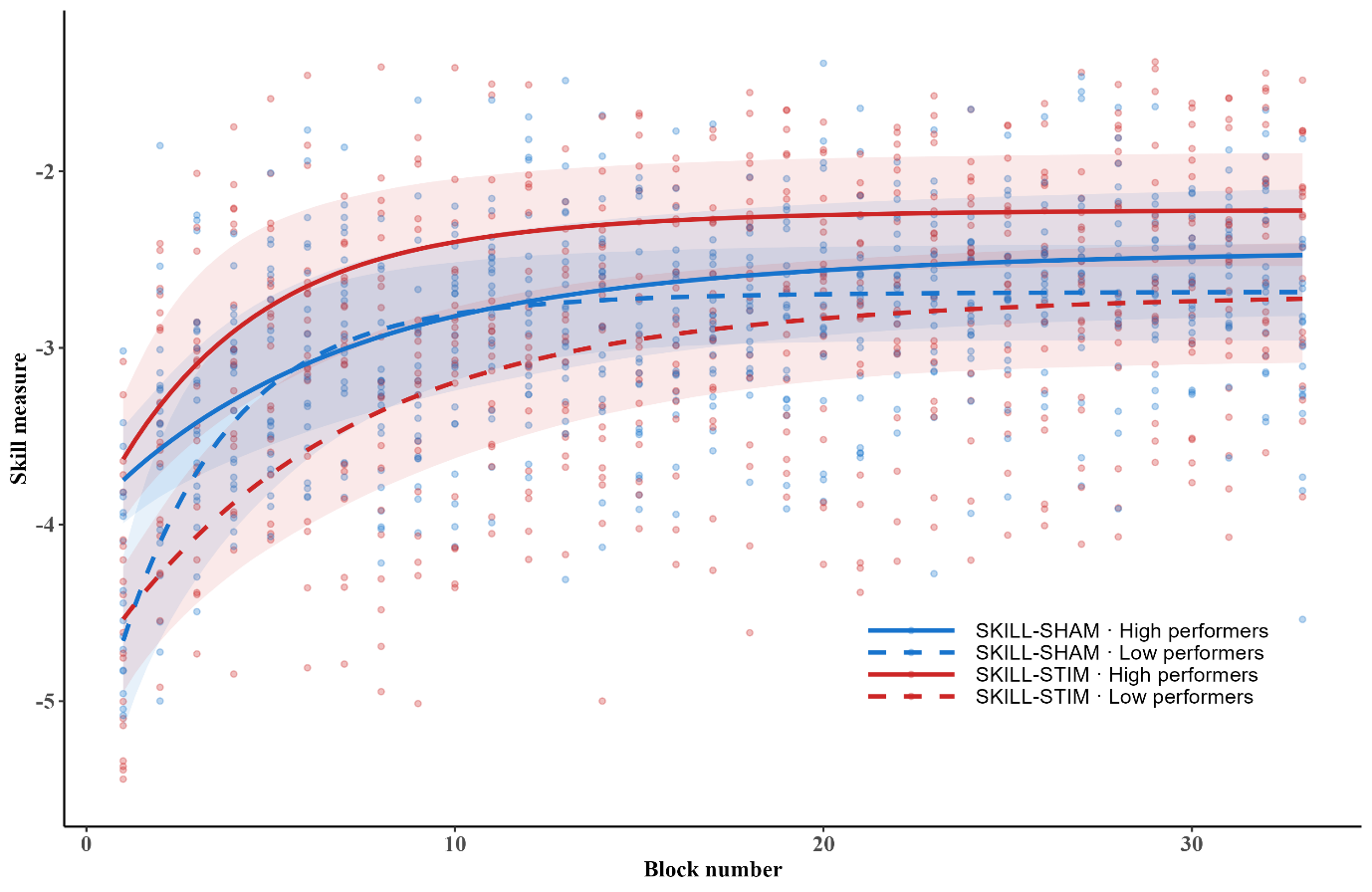

***Figure S3: High and low performers****. Grouped learning rates for high and low performers (classified from baseline performance within each group), showing the difference in learning rate across the intervention period; red color represents SKILL-STIM, blue color represents SKILL-SHAM, and grey represents CON.*

In exploratory subgroup analyses stratified by baseline performance, the estimated tDCS–sham difference in learning rate differed by subgroup. In high baseline performers, active stimulation was associated with a steeper learning rate than sham *(SKILL-STIM > SKILL-SHAM: −0.52 ± 0.52; 95% HDI [−1.38, 0.33], Evid. Ratio = 5.62, Post. Prob = 83%; ROPE = 3.78%)*. In low baseline performers, the pattern reversed *(SKILL-STIM > SKILL-SHAM: 0.79 ± 0.43; 95% HDI [0.08, 1.48], Evid. Ratio = 0.04, Post. Prob = 4%; ROPE = 1.47%)* (Fig. S3).
